# Mouse HP1γ regulates TRF1 expression and telomere stability

**DOI:** 10.1101/2022.11.24.517673

**Authors:** Emmanouil Stylianakis, Richard Festenstein, Jean-Baptiste Vannier

## Abstract

**Aims:** TElomeric Repeat-containing RNA are long non-coding RNAs generated from the telomeres. TERRAs are essential for the establishment of heterochromatin marks at telomeres, which serve for the binding of Heterochromatin Protein 1 (HP1), a protein family of epigenetic modifiers involved with chromatin compaction and gene silencing. While HP1γ is enriched on gene bodies of actively transcribed human and mouse genes, it is unclear if its transcriptional role is important for HP1γ function in telomere cohesion and telomere maintenance. We aimed to study the effect of mouse HP1γ on the transcription of telomere factors and molecules that can affect telomere maintenance.

**Main methods:** We investigated the telomere function of HP1γ by using deficient mouse embryonic fibroblasts (MEFs) deriving from 13.5 embryonic day embryos compared to their litter mate controls. We used gene expression analysis of HP1γ deficient MEFs and validated the molecular and mechanistic consequences of HP1γ loss by telomere FISH, immunofluorescence, RT-qPCR and DNA-RNA Immunoprecipitation (DRIP).

**Key findings:** Loss of HP1γ in primary MEFs leads to a downregulation of various telomere and telomere-accessory transcripts, including shelterin protein TRF1. Its downregulation is associated with increased telomere replication stress and DNA damage (γH2AX), effects more profound in females. We suggest that the source for the impaired telomere maintenance is a consequence of increased telomeric DNA-RNA hybrids and TERRAs arising at and from mouse chromosomes 18 and X.

**Significance:** Our results suggest an important transcriptional control by mouse HP1γ of various telomere factors including TRF1 protein and TERRAs that has profound consequences on telomere stability, with a potential sexually dimorphic nature.

## Introduction

Telomere stability requires the resolution of at least two important biological problems: the chromosome end-replication and the chromosome end-protection problems. The first refers to the gradual loss of genomic information through every cell cycle, owing mainly to incomplete lagging strand DNA synthesis [1-3] and resolved by the timely recruitment of telomerase reverse transcriptase [4]. The second problem refers to the quandary of the DNA damage repair machinery distinguishing the ends of linear chromosomes from DNA breaks resulting from damage caused by external (Ionising radiation) or internal sources (DNA replication stress).

Various mechanisms are in place for telomeres to avoid being recognized as DNA damage sites, including the actions of the shelterin complex [5]. Shelterin acts as a DNA damage response (DDR) inhibitor, masking telomeres from being sensed as double-strand breaks (DSBs) [6]. In addition to shelterin, vertebrate telomeres also contain heterochromatic histone marks that are characteristic of repressive or silent chromatin [7]. Studies in mice indicate a role for these epigenetic marks in telomere length maintenance and telomere recombination [8, 9]. One such mark, histone H3 trimethylated Lys 9 (H3K9Me3), provides a high-affinity binding site for heterochromatin protein 1 (HP1) [10, 11]. The HP1 protein family acts as elementary units of chromatin packaging and in mammals presents three isoforms: HP1α, HP1β and HP1γ [12]. The combination of high H3K9me3 density and HP1 occupancy leads to telomeric heterochromatinisation [13].

While all three HP1 isoforms are associated with heterochromatin [14], HP1γ is also found enriched on transcribing regions of the genome [15] and interacts with RNA polymerase II [16-18], suggesting for a role in transcription regulation.

RNA Polymerase II is responsible for the transcription at telomeres of noncoding RNAs termed TElomeric Repeat containing RNAs (TERRAs) [19-21]. TERRAs are crucial for the normal maintenance of telomeres by being an integral component of the telomeric chromatin [22]. TERRAs recruit Polycomb complex 2 to telomeres which facilitates the tri-methylation of lysine 27 on histone 3 (H3K27me3), leading to further establishment of H3K9me3 and the recruitment of HP1 [13].

A role for HP1 in telomere function is firmly established in Drosophila [23], suggested in mice [8, 9] and demonstrated in human cells for the establishment/maintenance of cohesion at telomeres [24]. However, the function of HP1γ at telomeres has never been investigated through the prism of its transcriptional function that the factor could have on TERRAs, while a negative feedback loop between TERRAs, H3K9me3 and RNA Polymerase II exists. Hence, the importance of the study to shed light on the cross-talk between the recruitment of HP1γ at telomeres by the axis TERRAs-Histone marks and HP1γ transcriptional regulation of TERRAs. In this study, we show that loss of HP1γ early in mouse development leads to the downregulation of various telomere and telomere-associated factors including TRF1 shelterin component, misregulation of TERRA expression and elevated levels of DNA-RNA hybrids at telomeres. Compromised telomere maintenance observed as telomere replication stress and telomere DNA damage induction is associated with elevated cellular senescence; highlighting the importance of HP1γ on telomere stability and cell physiology through its transcriptional regulation function.

## Materials and methods

### Animal handling and transgenic mice genotyping

Animals used in this study were maintained and handled according to the Imperial College London guidelines for Animal Research and the regulations of the British Home Office.

DNA was extracted from mouse ear punches or mouse embryo heads following the HotSHOT protocol [25]. The PCR samples were analysed with a 1.5 % agarose gel run at 100 V for 40 min. The list of primers and the PCR conditions can be found in tables S1-S4.

### Generation of mouse embryonic fibroblasts (MEFs) and cell culturing

HP1γ^+/+^ and HP1γ^-/-^ embryos were generated by crossing male and female HP1γ^+/-^ mice. For the generation of mouse embryonic fibroblasts (MEFs) E13.5 embryos were isolated. Embryo head and internal body organs were removed and the remaining body was finely minced in ice-cold 0.25% trypsin-EDTA (Sigma). The samples were incubated overnight at 4 °C and the following day, samples were incubated at 37 °C for 30 min and culture medium Dulbecco’s modified Eagle’s medium (DMEM) supplemented with 15% foetal bovine serum (FBS), 1% penicillin/streptomycin (Gibco) was added to the samples to deactivate trypsin. Vigorous resuspension allowed for isolation of the cells, which were subsequently cultured at 37 °C and 5% (v/v) CO_2_.

### Senescence-associated β-galactosidase assay (SA-β-gal)

Cells were washed twice with 1x phosphate-buffered saline (PBS) before being fixed with 0.5% Glutaraldehyde (Sigma-Aldrich, G7776) in 1x PBS for 15 min, followed by 2 washes with 1x PBS. X-gal solution (1 mg/ml X-gal, 5 mM potassium ferrocyanide, 5 mM potassium ferricyanide,1 mM MgCl2 in 1x PBS) was used to stain the cells by incubating at 37 °C for 16 h. After staining, the cells were washed with 1x PBS and the proportion of cells with SA-β-gal activity was quantified using the Olympus CKX41 microscope.

### Quantitative-FISH analysis (Q-FISH)

For metaphase spread preparation, MEFs were incubated for 4 h with 10 ng/ml colcemid (Roche, 10295892001). Cells were then collected and incubated for 15 min at 37 °C, in hypotonic buffer (KCl 75 mM). Fixation was performed with ethanol: glacial acetic acid (3:1, v/v), followed by three washes with the same fixative. The metaphase suspensions were subsequently dropped on glass slides.

Q-FISH was performed as previously described [26]. Briefly, metaphase spreads were fixed with 4% formaldehyde for 2 min, washed three times in 1X PBS, and treated with pepsin (1 mg/ml in 0.05 M citric acid pH=2) for 10 min at 37 °C. They were post-fixed for 2 min with 4% formaldehyde, washed three times with 1X PBS and incubated in increasing ethanol concentration baths. Each slide was then covered with hybridising solution containing Cy3-O-O-(CCCTAA)_3_ probe (PNA bio) in 70% formamide, 10 mM Tris pH=7.4 and 1% blocking reagent (Roche, 11096176001). This step was followed by denaturation for 3 min at 80 °C. Hybridisation was performed for 2 h at room temperature and the slides were washed twice for 15 min in 70% formamide, 20 mM Tris pH=7.4. Three 5 min washes in 50 mM Tris pH 7.4, 150 mM NaCl, 0.05% Tween-20 followed and the slides were dehydrated in successive ethanol baths and air-dried. Slides were mounted with ProLong Gold antifade reagent containing DAPI (Invitrogen) and images were captured with Zeiss microscope using Carl Zeiss software. Telomeric signal was quantified using the ImageJ FIJI software.

### Immunofluorescence (IF)-FISH

MEFs seeded on culture slides, were permeabilised with Triton X-100 buffer (50 mM NaCl, 3 mM MgCl2, 20 mM Tris pH=8, 0.5% Triton X-100, 300 mM sucrose), fixed for 15 min in fixative solution (4% formaldehyde, 2% sucrose) and washed three times with 1x PBS. Another 10 min permeabilisation followed, slides were washed once with 1x PBS and were incubated for 30 min with blocking buffer (10% goat or donkey serum (Stratech Scientific Ltd) in 1X PBS) at 37 °C. Then, the primary antibody (anti-γH2AX, Millipore, 05-636, 1:500; anti-53BP1, Thermo Fisher Scientific, 1:400) was added in blocking buffer and incubated for 1 hour at 37 °C. Three washes with 1X PBS, followed and the slides were incubated with secondary antibody (1:400 in blocking buffer, donkey a-rabbit Alexa 488 antibody, Invitrogen A-21206; or goat a-mouse Alexa488, Invitrogen A-11001) for 30 min at 37 °C. Slides were then post-fixed for 10 min using 4% formaldehyde, 2% sucrose and washed three times with 1x PBS for 3 min. FISH was then performed as in metaphases, using the Cy3-O-O-(CCCTAA)_3_ probe.

Images were captured with Zeiss microscope using Carl Zeiss software and quantification was performed using the CellProfiler 3.1.9 software.

### Northern blot

RNA extraction was performed using the RNeasy Mini Kit (Qiagen, 74104) and DNA was eliminated by on-column treatment with DNase I (Qiagen, 79254). 10 µg of total RNA was denatured for 10 min at 65°C in 1X MOPS (0.2M MOPS, 50 mM NaOAc, 10 mM EDTA, RNase-free water) with 50% formamide, and 2.2M formaldehyde, followed by a 5 min incubation on ice. 10X dye buffer (50% Glycerol, 0.3% Bromophenol Blue, 4 mg/ml Ethidium Bromide) was added to the samples, which were then run on a formaldehyde agarose gel (0.8% agarose, 1X MOPS, 6.5% formaldehyde) at 5 V/cm in 1X MOPS buffer. The gel was subsequently washed twice with RNase-free water and three times with 20X SSC, before transferring the RNA to an Amersham Hybond N+ membrane (Cytiva) using a neutral transfer in 20X SSC. The membrane was UV-crosslinked (Stratalinker, 2000 kJ) and baked for 45 min at 80 °C, followed by hybridisation with a ^32^-P labelled TAA(CCCTAA)_4_ probe using UltraHyb-Oligo solution (ThermoFisher #AM8663) at 42 °C overnight. The next day, the membrane was washed with 2X SSC, 0.1% SDS for 10 min at room temperature, 5 min with 0.2X SSC, 0.1% SDS at 42 °C and for 5 min with 2X SSC, 0.1% SDS at room temperature. Signal was detected with phosphor-imager (Amersham Biosciences).

### Telomere restriction fragment (TRF) analysis

DNA was isolated from cells, using the Hirt buffer (10 mM Tris pH=7.6, 100 mM NaCl, 10 mM EDTA) with the addition of 0.5% SDS and 200 μg/ml of RNase A (Roche, 10109169001). The samples were heated at 37 °C for 60 min, followed by the addition of 150 μg/ml Proteinase K and further incubating the samples at 55 °C overnight. 5 μg of DNA was digested with HinfI and RsaI restriction enzymes (New England BioLabs, 50U each) at 37 °C overnight. Digested DNA was purified with phenol-chloroform and 5μg were loaded on a 0.8 % agarose gel and run at 40 V for 20 h. DNA was then depurinated for 15 min with 250 mM HCl, denatured for 30 min in 500 mM NaOH,1.5 M NaCl, neutralized for 30 min in 1.5 M NaCl, 500 mM Tris PH=7.5 and transferred overnight on an Amersham Hybond N+ membrane with 20x SSC. DNA was UV-crosslinked (7000 kJ) and the membrane was neutralized with 2x SSC. The membrane was hybridized with a TAA(CCCTAA)_4_ probe which was conjugated with digoxigenin (DIG oligonucleotide 3⍰-end labelling kit, Roche) and signal was revealed using the anti-DIG-alkaline phosphatase antibodies (Roche) and CDP-Star (Roche), following the manufacturer’s instructions. Images were captured using an Amersham Imager 680.

### RNA dot blot

RNA extraction was carried out similar to Northern blotting. 2 µg of the extracted RNA were treated with RNase A (500 μg/ml) for 3 h at 37 °C to serve as negative control. RNase A treated and RNase A non-treated samples (2 μg) were denatured in 0.2 M NaOH by heating at 65°C for 10 min, incubated 5 min on ice and spotted on an Amersham Hybond N+ membrane. RNA was then UV-crosslinked (2000 kJ) and the membrane was baked for 45 min at 80°C. Hybridisation of the membrane followed similarly to the TRF assay using the DIG-labeled telomeric C-rich probe. The membrane was imaged using the Amersham Imager 680 and analysed using the Image Studio Lite software. The membrane was then stripped and re-probed with an 18s rRNA targeting probe (5’-CCATCCAATCGGTAGTAGCG-3’, Sigma) which was used for signal normalisation.

### TERRA RT-qPCR

RT-qPCR for TERRAs was performed as previously described [27] with some modifications. RNA was extracted similar to Northern blotting with two additional DNase I digestions. 1U DNase I (Roche) per μg of RNA was used followed by an on-column treatment with DNase I (Qiagen) according to the manufacturer’s protocol. 3 μg of total RNA was reverse transcribed using 200 U of SuperScript III Reverse Transcriptase (Invitrogen) with either random hexamers (Invitrogen, N8080127) or TERRA specific oligonucleotides (CCCTAA)_4_ [28]. qPCR reactions were performed using Power SYBR Green PCR Master Mix (Applied Biosystems, 4368708) and the Bio-Rad CFX96 system with the following parameters: 40 cycles of 15s of denaturation at 95 °C followed by 1 min of annealing and extension at 60 °C. RT-qPCR primers are listed in table S3. The 2_t_^-ΔΔCt^ method was employed for relative TERRA quantification, using the β-actin housekeeping gene for normalisation.

### DNA-RNA Immunoprecipitation (DRIP)

DRIP was performed as previously described [29, 30], with the following modifications. Genomic DNA was extracted from MEFs using phenol-chloroform. DNA was digested at 37 °C for 16 h with the following restriction enzymes: SspI, EcoRI, HindIII, XhoI and BsrGI (New England BioLabs, 44 U each). Digested nucleic acids were EtOH-precipitated with 1.5 μl of glycogen (Thermo Fisher Scientific, R0561) and resuspended in 50 μl of 1x TE buffer. 6 μg of digested nucleic acids were treated overnight at 37 °C with or without 10 μl of RNase H (New England BioLabs, M029L) in 1x RNase H buffer and 1/10^th^ of the samples was saved as input. RNase H treated and non-treated samples were incubated with 10 μl of the S9.6 antibody (Kerafast ENH001) for 16 h at 4 °C, followed by an incubation of 100 μl of a 2:1 (A:G) mixture of Dynabeads Protein A and G (Invitrogen, 10001D and 10004D) for 2h at 4 °C. The precipitated samples were eluted in 300 μl of elution buffer (10 mM EDTA PH= 8, 50 mM Tris PH=8, 0.5% SDS) and treated with 7 μl of proteinase K (24 mg/mL, P4850) for 45 min at 55 °C. A treatment with 50 μg/ml RNase A (Roche, 10109169001) for 1 h at 37 °C plus 1 h at 65 °C followed and cleaned samples were resuspended in 100 μl of 1X TE buffer. DNA-RNA hybrids were detected using at the indicated subtelomeric regions with the corresponding qPCR primers listed in table S3 [28, 31]. qPCR was performed with the same parameters employed for the TERRA RT-qPCR and percentage of signal to input was calculated.

### Protein extraction and western blotting (WB)

Cell pellets were incubated for 10 min on ice in lysis buffer (25 mM Tris PH=8, 40 mM NaCl, 2 mM MgCl_2_, 0.05% SDS) supplemented with 100 units/ml Benzonase (Sigma-Aldrich, E1014-25KU) and cOmplete, EDTA-free Protease Inhibitor Cocktail (Roche). The samples were lysed by being forced ten times through a 25G needle and incubated on ice for a further 10 min. Proteins were quantitated using the Pierce BCA Protein Assay kit (Thermo Fisher Scientific, #23225) and 30 μg from each sample were denatured for 5 min at 100 °C with the addition of 4x Laemmli buffer (50 mM Tris pH=6.8, 2% SDS, 100 mM DTT, 10% glycerol, 0.1% bromophenol blue (w/v)). Proteins were resolved in 4–12% Bis-Tris gels (Invitrogen) at 100 V for 2.5 h and then transferred to a nitrocellulose membrane (Amersham Protran 0.2 µm NC), employing a wet transfer apparatus (Bio-Rad) at 90 V for 2h at 4 °C. After blocking with 5% non-fat milk in 1x PBS, 0.1% Tween (PBST) for 1 h at room temperature the membranes were probed overnight with primary antibodies mouse anti-HP1γ (1:2,500) (Thermo Fisher Scientific #MA3-054), rabbit anti-TRF1 (gift from Titia de Lange) and mouse anti-α-tubulin (1: 5,000) (Sigma-Aldrich #T6199) at 4 °C. The membranes were washed three times with 1x PBST for 5 min and incubated with secondary antibodies anti-mouse-HRP (1:10,000) (Agilent Dako, #P0447) and anti-rabbit-HRP (1:10,000) (Agilent Dako, #P0217) for 1 h at room temperature, followed by 1x PBST washes. Finally, the signal was visualized using ECL Western blotting reagent (Sigma-Aldrich, RPN2106) and exposed with the Amersham Imager 680 (GE Healthcare).

### Differential gene expression analysis and statistical analysis

RNA-sequencing data (Preprint BioRxiv: https://doi.org/10.1101/563940) was examined with the DESeq2 Bioconductor package [32]. Raw p-values were adjusted for multiple testing using the Benjamini-Hochberg procedure. Statistical analysis for all other experimental procedures was performed using GraphPad Prism. Error bars, number of replicates (n) and statistical methods are indicated in figure legends. Biological replicates for all experiments were based on embryos from independent litters.

## Results

### Mouse HP1γ regulates an extensive list of factors involved directly and indirectly in telomere maintenance without affecting telomere length

Given the recruitment of HP1 at telomeres for heterochromatin establishment [13], we utilised an HP1γ knockout mouse model [33], to elucidate the role of this isoform on telomere maintenance. HP1γ deficiency leads to neonatal lethality [34], however, mice with one mutated CBX3 allele (HP1γ^+/-^) present no apparent phenotype and develop normally into adulthood. By breeding HP1γ^+/-^ mice, we were able to isolate mouse embryonic fibroblasts (MEFs) deriving from embryonic day 13.5 (E13.5) embryos (Figure 1A and S1A). In homozygous HP1γ knockout (HP1γ^-/-^) MEFs, HP1γ RNA and protein levels are undetectable compared to the proficient MEFs (HP1γ^+/+^) derived from littermates (Fig 1B & S1B).

**Figure 1:**
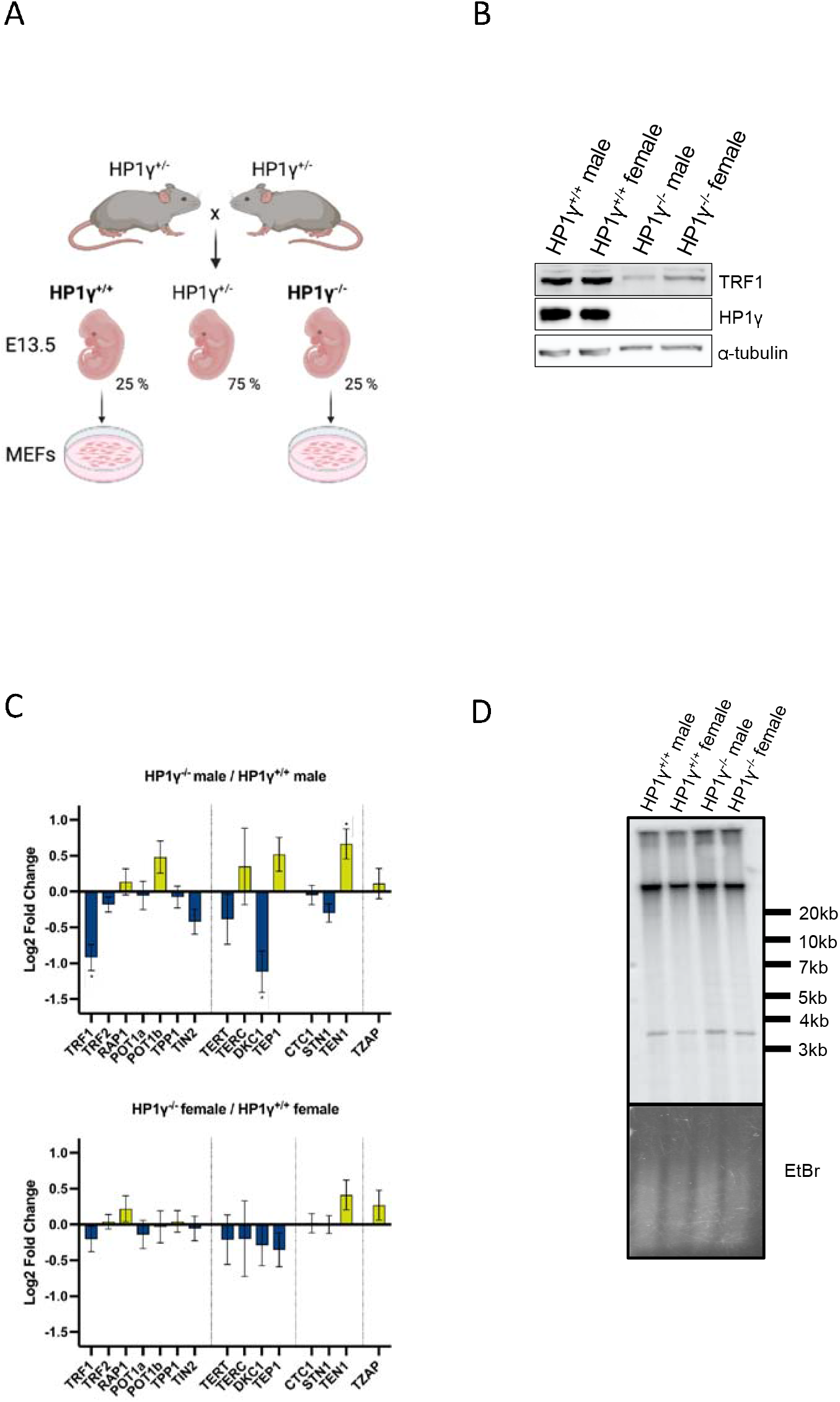
(A) Schematic of crossing strategy to generate homozygous HP1γ knockout (HP1γ^-/-^) embryos. Inter-breeding of heterozygous HP1γ (HP1γ^+/-^) mice results in a 25% chance of wild-type (HP1γ^+/+^) embryos, 25% chance of HP1γ^-/-^ and 75% chance of HP1γ^+/-^ embryos. Mouse embryonic fibroblasts (MEFs) are generated from embryonic day 13.5 (E13.5) embryos. (B) Protein expression of HP1γ, TRF1 and α-tubulin (loading control) by western blot. 30 μg of proteins were loaded for each condition. (C) Changes in expression of factors that are necessary for telomere stability upon HP1γ depletion. Analysis of mouse shelterin genes (TRF1, TRF2, RAP1, POT1a, POT1b, TTP1, TIN2), telomerase-associated genes (TERT, TERC, DKC1, TEP1) CST complex (CTC1, STN1, TEN1) and TZAP. Bars represent the Mean log2 fold changes ± SEM of three biological replicates for each genotype (Preprint BioRxiv: https://doi.org/10.1101/563940). * padj < 0.05. (D) Telomere Restriction Fragments (TRF) analysis by southern blot. The blot was revealed with a DIG-Tel-C-rich probe (top). Ethidium bromide (EtBr) staining (bottom) is used as loading control.

Since HP1γ is capable of transcriptional regulation [15] and telomere maintenance [24], we decided to look for the possible transcriptional regulation of factors that are necessary for telomere stability. We analysed publicly available transcriptomic data of MEFs lacking HP1γ (Preprint BioRxiv: https://doi.org/10.1101/563940) to test this hypothesis. Especially in males, the absence of HP1γ leads to a widespread dysregulation of factors involved directly with telomere maintenance, including shelterin TRF1 and telomerase associated Dyskerin (DKC1; Figure 1C) but also telomere-associated helicases RTEL1, BLM, FANCM and DNA repair/signalling proteins including RAD51 and its paralogs (Figure S1C). We further confirmed that downregulation of TRF1 transcripts observed in HP1γ^-/-^ cells was conserved at the protein level by Western Blotting for both males and females HP1γ^-/-^ MEFs (Figure 1B). TRF1 is an essential telomere factor facilitating telomere replication [35, 36] and averts mouse telomere chromatin remodelling [37].

Despite the transcriptional effect of HP1γ on some telomerase-associated and telomere factors, the average telomere length, measured by southern blotting, remained unchanged among the different genotypes and sexually unbiased (Figure 1D). The majority of telomeres migrated into a strong band above the 20 kb mark and the smearing of shorter telomeres remained faint in all conditions. HP1γ loss has no major noticeable effect on telomere length. This is also consistent with previous reports about the absence of effect on telomere length when mouse TRF1 is downregulated [35, 36].

### HP1γ regulates TERRA levels and telomeric DNA-RNA hybrids

The role of HP1γ on transcription regulation is still an open question, while some suggest an activating role [17] and others report a silencing effect [38]. Remarkably, we observed a significant reduction in TRF1 proteins in male and female HP1γ^-/-^ cells, to levels of a knockdown (Figure 1C). Some of the known consequences of mTRF1 dysregulation are increased TERRA levels and telomere replication stress (measured as fragile telomeres; [37]). First, we utilised previously generated RNA-sequencing data from captured mouse polyadenylated RNA (Preprint BioRxiv: https://doi.org/10.1101/563940) to align an *in-silico* TERRA probe to the sequencing reads. This allowed us to identify the polyadenylated TERRA transcripts. Our analysis revealed no major differences regarding this fraction of TERRAs among the wild-type sexes. Interestingly, upon loss of HP1γ, the polyadenylated TERRA transcripts almost double in males (1.8-fold difference) and more than triple in females (3.5-fold difference; Figure S2A).

To further characterise the effects of HP1γ loss on TERRA levels, RNA dot blot and Northern blot analyses were performed using total RNA from E13.5 MEFs. In agreement with the RNA-sequencing analysis, RNA dot blot revealed an increase of RNA molecules containing telomeric repeats in both male and female HP1γ^-/-^ conditions (Figure S2B). More precisely, the Northern blot showed a sharp increase of high molecular weight TERRAs (over 9kb), particularly evident in HP1γ^-/-^ females (Figure 2A).

**Figure 2:**
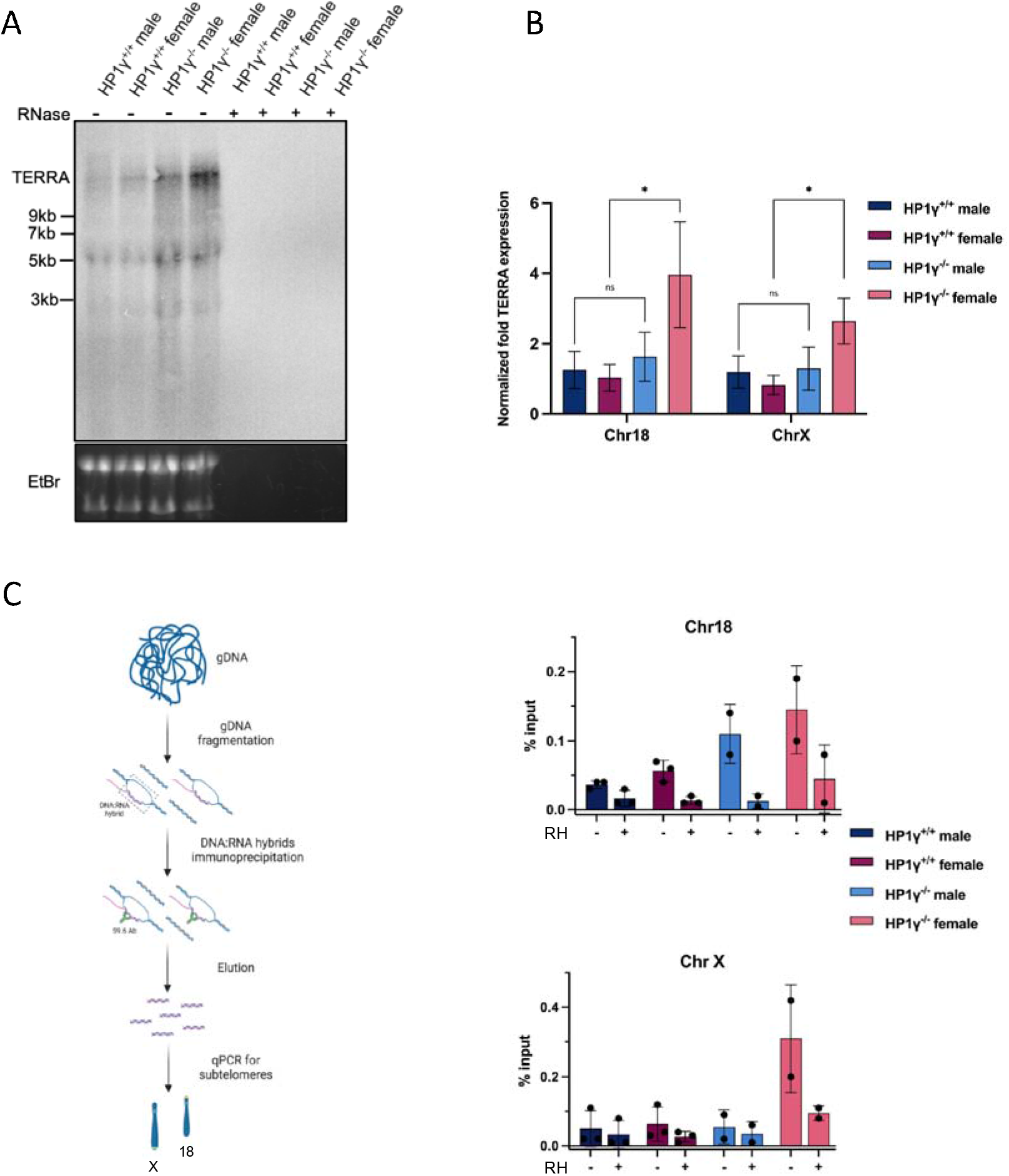
(A) Northern blotting of total RNA (10 μg) using a ^32^P-labelled telC probe for hybridisation and detection of telomeric TERRAs. RNAse A treatment is used to certify specific RNA signal (+). Before transfer onto Nylon+ membrane, nucleic acids inside the gel are detected following incubation in Ethidium Bromide (EtBr) buffer, imaged and act as a loading control. (B) RT-qPCR of total MEFs RNA (3 μg). The graphs represent the average value from three biological and three technical replicates for each sample after normalisation to β-actin. Error bars represent the SD. Statistical significance was tested with one-way ANOVA and Tukey’s post-hoc test, * p<0.05 ns=non-significant. (C) Left: Outline of the DRIP experiment. Briefly, genomic DNA (gDNA) is isolated and fragmented by a combination of restriction enzymes. DNA:RNA hybrids are immunoprecipitated with the S9.6 antibody, the fragments are isolated and quantification of site-specific DNA:RNA hybrids is performed by qPCR. Right: DRIP RT-qPCR quantification for chromosomes 18 and X subtelomeric regions. Bars represent the mean ± SD for the percentage of DNA:RNA hybrids in respect to input for three biological replicates of HP1γ^+/+^ male and HP1γ^+/+^ female MEFs and two biological replicates of HP1γ^-/-^ male and HP1γ^-/-^ females MEFs. RH: RNase H treatment act as control testing specificity of the signal towards DNA:RNA hybrids.

In mice, the vast majority of TERRAs arise from the subtelomere of chromosome 18, while a smaller fraction may arise from other subtelomeric regions including those on chromosome 9, 10 and X [28, 39, 40]. Utilizing specific primers for the subtelomeres of chromosomes 9, 10, 18 and X, we observed by RT-qPCR that TERRAs arising from chromosomes 18 and X are significantly increased in female HP1γ^-/-^ MEFs (Figure 2B). However, we could not detect a significant variation of TERRAs arising from chromosome 18 and X subtelomeres for the male HP1γ^-/-^ MEFs. TERRAs formation from chromosome 9 and 10 remained at undetectable levels, irrespective of the genotypes (data not shown).

Due to the complementarity of TERRA molecules with telomeric DNA, TERRAs can hybridize to the exposed C-rich lagging strand during DNA replication and generate DNA-RNA hybrids at telomeres, which in turn can give rise to R-loops by displacing the G-rich strand [41]. While these structures appear to be important for physiological telomere homeostasis [41-43], persistent DNA-RNA hybrids can cause replication fork stalling and hence replication stress [44-47].

We examined whether the increased TERRA levels observed in HP1γ depleted cells lead to more telomeric R-loops by performing DNA-RNA immunoprecipitation (DRIP) experiments (Figure 2C). During DRIP, the S9.6 monoclonal antibody is employed to specifically immunoprecipitate DNA-RNA hybrids [48, 49]. Characterisation of the localisation of the formation of these structures can be achieved with subsequent qPCRs. Since dysregulation of TERRA expression was mainly observed on the end of chromosomes 18 and X, we analysed the abundance of DNA-RNA hybrids at these two specific chromosome-ends. Telomeric DNA-RNA hybrids signal increased upon HP1γ loss at the ends of chromosomes 18 and X, especially in female knockout MEFs, and was suppressed by RNaseH1 treatment (Figure 2C). This indicates that HP1γ is essential to prevent increased TERRA levels that can form excessive telomeric DNA-RNA hybrids at chromosomes 18 and X telomeres, which could be mediated via an indirect regulation of TRF1. Interestingly, female embryos and cells might be more sensitive to such dysregulation in the absence of HP1γ.

### HP1γ-TRF1 axis protects telomeres from telomere replication stress, DNA damage and senescence

Given the downregulation of TRF1, increased TERRA expression and elevated levels of telomeric DNA-RNA hybrids in the absence of HP1γ, we would expect a defect in telomere integrity. Fragile telomeres are characterised by aberrant signal that is spatially separated from the chromatid end of metaphasic chromosomes [35]. We used Quantitative-fluorescence in situ hybridisation (Q-FISH) experiments of metaphase spreads and observed no significant differences in the stability of telomeres between wild-type males and females (Figure 3A). However, comparison of HP1γ^+/+^ and HP1γ^-/-^ telomeres revealed a significant higher incidence of fragility upon loss of HP1γ. The effect is stronger in females where 17% of the HP1γ^-/-^ chromosomes have fragile telomeres compared to 9% in HP1γ^+/+^ females, 11% HP1γ^+/+^ males and 14% in HP1γ^-/-^ males (Figure 3A), consistent with the sexual bias observed for R-loops proportions. Telomere fragility is the main telomere defect in HP1γ^-/-^ MEFs as neither telomere fusions nor telomere loss were observed (Figure S3A), which is also consistent with a TRF1-like effect in these cells [37]. Together the results suggest that the increased telomeric fragility observed in females arises not only due to the TRF1 downregulation that occurs in both sexes, but also due to the increased TERRAs and R-loops particularly evident in females HP1γ^-/-^. Telomere fragility is a marker of replication stress that is associated with increased telomere DNA damage [35, 36, 50]. Therefore, we decided to look for a DNA damage response following HP1γ using immunofluorescence of the phosphorylated histone variant H2AX (γH2AX). HP1γ deletion leads not only to a nuclear-wide increase of γH2AX immunofluorescence signal, but also to more telomere dysfunction-induced foci (TIFs) (Figure 3B). Unlike γH2AX, 53BP1 (DSB marker) levels in cells and at telomeres are not increased in HP1γ^-/-^ conditions compared to HP1γ^+/+^ MEFs (Figure S3B).

**Figure 3:**
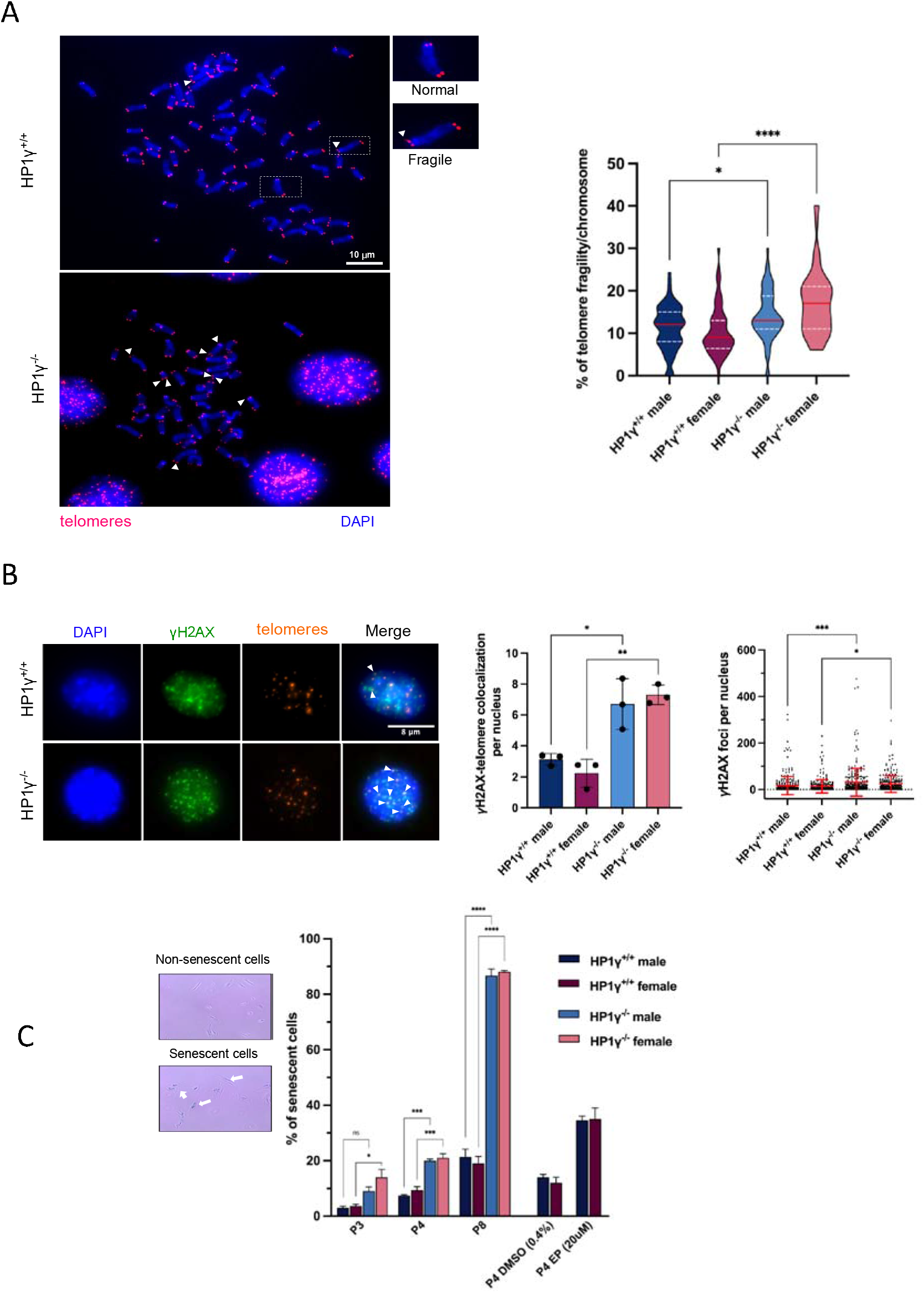
(A) Left: Representative images of mouse metaphasic chromosomes (DAPI, blue), hybridized with a telomere-specific probe, conjugated with a Cy3 fluorophore (red). The arrows indicate distorted signal (fragile telomeric site). Right: Graphical quantification of the percentage of telomere fragility per chromosome. Violin plots represent the combined analysis of 60 metaphases from three independent biological replicates (20 metaphases per biological replicate). Median is coloured in red and quartiles in white. Statistical significance was tested with one-way ANOVA and Tukey’s post-hoc test, * p<0.05, **** p<0.0001. (B) Left: Representative image of IF-FISH showing co-localisation of telomeres (red) with γH2AX (green) in HP1γ^+/+^ and HP1γ^-/-^ MEFs nuclei (DAPI, blue). Right: Quantification of γH2AX co-localisation to telomeres per nucleus (n = 300 nuclei) and total γH2AX foci per nucleus. Data are represented as mean ± SD from three independent biological replicates. Statistical significance was tested with one-way ANOVA and Tukey’s *post-hoc* test, * p<0.05, ** p<0.01, *** p<0.001. (C) Representative images of SA-β-gal staining for senescent (light blue coloured cells, white arrows) and non-senescent cells. Bars represent the mean ± SEM from three biological replicates (at least 100 cells analysed per replicate) of the percentage of senescent cells at Passage 3, 4 and 8 of primary MEFs that are proficient and deficient for HP1γ. Two biological replicates of Passage 4 HP1γ^+/+^ cells were also treated with either DMSO (0.4%) or Etoposide (20uM) as negative and positive controls for senescence, respectively. Statistical significance was tested with one-way ANOVA and Tukey’s post-hoc test, * p<0.05, ** p<0.01, *** p<0.001, **** p<0.0001.

Compromising the telomeric structure and inducing DNA damage are sources of cellular senescence, a state where there is loss of cell proliferative capacity, despite continued metabolic activity and viability [51]. We noticed a rapid change in size and morphology of MEFs lacking HP1γ, with cells becoming larger and more spindle-shaped, suggestive of a senescent state [52]. By testing for β-galactosidase (β-gal) activity, one of the most commonly used senescence biomarkers [53], we addressed whether earlier onset of senescence arises due to HP1γ loss and subsequent telomere maintenance defects. We observed that less than 10% of the HP1γ wild-type (HP1γ^+/+^) male and female MEFs were senescent at passaging 3 and 4 (P3 and P4; Figure 3C). Interestingly, at P3 more than 15% of the HP1γ^-/-^ female cells are senescent and increases to 20% at P4 for both sexes and by P8, a drastic surge in senescence is observed, where almost 90% of HP1γ^-/-^ cells are senescent for both sexes. As a control, we used a 20µM etoposide treatment for 16 hours on wild-type MEFs to ensure that β-gal assay was working properly. Indeed, we could observe a 3-fold increase of senescence cells. To conclude, loss of HP1γ leads to telomeric DNA damage induction and earlier onset of senescence.

## Discussion

Telomere stability is largely influenced by the chromatin state. While recent studies show that human telomeres display lower than expected levels of H3K9me3 [54, 55], mouse telomeres are naturally enriched for this heterochromatic mark [8], and have a highly compacted chromatin state [56]. Histone methyltransferase SETDB1, known interactor of HP1 [57], is responsible for the establishment of H3K9me3 on mouse telomeres [58], and loss of other H3K9me3-affiliated HMTs like SUV39H1/H2 results in defective telomeres with increased length [8]. HP1γ is a member of the HP1 proteins, a family of proteins necessary for the establishment and propagation of heterochromatin [12], a chromatin state long thought to be transcriptionally silent. At the same time, HP1γ is an essential factor for gene expression [15, 59], enriched on gene bodies of actively transcribed genes [16, 18]. HP1γ offers a dilemma of functions that makes it particularly interesting in the context of telomere maintenance where transcription of telomeric lncRNA molecules TERRAs is cell cycle regulated and essential for the establishment of heterochromatic marks [19]. To understand this quandary, we decided to take advantage of the known recruitment of HP1γ at telomeres via heterochromatin marks, including H3K9me3 [13] and its involvement with telomere cohesion and maintenance [24] to answer the question: Is HP1γ control of telomere maintenance at early mouse development mediated by transcriptional regulation?

Here, we report that HP1γ regulates the expression of many telomere and telomere-accessory factors that are related to facilitating telomere replication and suppressors of telomere fragility [60]. This corresponds to DKC1 that encodes dyskerin involved in the maturation of TERC [61], RTEL1 helicase that facilitates DNA replication through telomeric DNA secondary structures [50] or the BLM helicase that resolves G4 structures [62], many other DNA repair and recombination factors and including TRF1 shelterin factor, that are all transcriptionally downregulated in the absence of HP1γ, especially in males. Mis-regulation of most of these factors have been associated with increased replication stress at telomeres [60], including TRF1 that results in the so-called telomere fragility [35, 36, 60].

HP1γ’s interaction with TIN2 has been shown to be necessary for the establishment of cohesion at human telomeres in S phase [24]. The lower levels of TRF1 in the absence of HP1γ could further disrupt shelterin formation and implicate an indirect role of HP1γ in telomere cohesion through its transcriptional role. Since most of the telomere-associated genes are downregulated upon HP1γ knockout, it is likely that for these genes, HP1γ acts as a transcriptional activator.

Telomeres were long considered silent, but the discovery of TERRAs, transcripts from these regions, broke the dogma [63, 64]. HP1γ appears to affect TERRA functionality by influencing their numbers. Total TERRA numbers but also the polyadenylated TERRA fraction, which comprises the majority of TERRA molecules in mice [28] are elevated in the absence of HP1γ. This increase in TERRA signal is not a consequence of altered telomere length. Indeed, longer telomeres could act as extended substrates for TERRA transcription [65] and shorter telomeres could lead to a lower number of transcription repressors and in turn increased TERRA expression as seen in yeast, [66, 67]. In our study, the unchanged telomeric length in HP1γ^-/-^ MEFs supports a direct regulation HP1y on the expression of mouse TERRAs.

Also, the higher levels of TERRAs without changes in their length (from the Northern blot) support that HP1γ’s role is not with TERRAs transcriptional termination nor with TERRA processing by splicing regulation [68, 69]. Both would have led to differences in the length of TERRA transcripts because of uncontrolled transcriptional termination at multiple sites within the telomeric tract and/or differential processing of their polyadenylation or because of miss-splicing of transcripts. If the splicing variants were unstable, they would have been degraded at a faster rate [63] and the total TERRA levels would have been lower in HP1γ^-/-^ MEFs.

Another possibility of how HP1γ may contribute to the suppression of TERRAs is by promoting the formation of a locally condensed chromatin structure. Since HP1γ physically interacts with H3K9 HMTs [57, 70, 71], their recruitment can induce further methylation of H3K9 at the target locus which in turn can provide more binding sites for the HP1 isoforms. Dense HP1 binding and dimerisation can result in the re-configuration of the histone core with the normally buried residues now exposed and able to participate in weak multivalent interactions, promoting the formation of phase-separated liquid condensates and tight crosslinking of nucleosomes [72]. While HP1α has been shown to form liquid droplets on its own [73, 74], HP1γ heterodimerisation with other HP1 proteins [75, 76] may result in LLPS-driven exclusion of transcription factors and ultimately silencing of transcription. Further work is necessary to elucidate this aspect.

An important portion of the total mouse TERRAs arise from chromosome X [28, 77] and since females have two X chromosomes, one might expect that expression of X-linked TERRAs would be double in females. However, the RT-qPCR analysis argues that both sexes are showing comparable levels in wild-type conditions. The inactivation of one of the two X chromosomes could be acting as a dosage compensation mechanism also for these telomeric long-non coding RNAs.

HP1γ is potentially influencing telomere chromatin at least at two levels. Via its direct presence at telomeres [24] and via this newly described transcriptional effect on TRF1 regulation. Since one of the most notable effects of mouse TRF1 loss on telomeres is replication stress, the observation in HP1γ^-/-^ MEFs of increased fragile telomeres to similar levels to TRF1^-/-^ MEFs was expected [35, 36]. Fragile sites represent genomic regions where replication forks stall and/or collapse with their repair requiring breakage and re-establishment of functional DNA synthesis [78]. Problems during this process result in fragile sites being hotspots for chromosomal rearrangements [79].

Microscopy experiments followed to further investigate the potential DNA damage caused by the deletion of HP1γ by targeting proteins like γH2AX and 53BP1, early markers of DNA damage [80]. Our analysis revealed that upon depletion of HP1γ, γH2AX and 53BP1 levels rise in the nucleus, with a significant enrichment of γH2AX at telomeric sites (TIFs). The absence of 53BP1 at telomeres and the lack of telomeric fusions in HP1γ^-/-^ MEFs suggest for an accumulation of Single Strand Breaks (SSBs) at telomeres [81]. Telomeric SSBs would likely arise from telomere replication fork stalling/collapsing and reflected as fragility in the absence of TRF1. Unresolved SSBs would lead to detrimental DSBs, rendering the selection of the damage repair pathway critical for the maintenance of genomic stability [82]. The lack of 53BP1 TIFs and decreased TRF1 levels in HP1γ^-/-^ MEFs, would indicate for HR to be the repair pathway of choice, where the repair machinery is recruited following telomeric replication stress [83]. As BLM has been shown to drive BIR-mediated HR at telomeres [84], its downregulation in HP1γ depleted cells argues against the activity of this pathway and could potentially exacerbate the fragile phenotype [85].

Several studies have attributed a repressive role to TRF1 regarding TERRA regulation with several reports showing that loss of TRF1 lead to an upregulation of TERRAs in human and mouse cells [19, 37, 86, 87]. Therefore, the combination effect of HP1γ depletion, TRF1 downregulation and accumulation of TERRAs at telomeres were expected to increase the number of telomeric R-loops in HP1γ^-/-^ conditions, which would be the most propitious source for contributing to the telomere fragility and TIFs [41, 88]. R-loops are dynamic structures linked to ongoing transcription and known to form in response to transcriptional or replicative induced torsional stress in double-stranded DNA [89]. While TERRA transcription is programmed to not overlap with replication through the cell cycle in human cells, with highest TERRA levels in early G1 and at their lowest in late S phase [19] and since TERRAs can form telomeric R-loops post-transcriptionally and *in trans* [86], it is unclear at which stage of the cell cycle the TERRA-mediated R-loops on mouse chromosomes 18 and X are formed. Despite the importance of these structures for physiological telomere homeostasis [41-43], persistent R-loops can cause replication fork stalling and hence replication stress [44-46]. The formation and dissolution but also the existence of various backup pathways of degradation and repair ensures that R-loops biology are well regulated. For instance; in Immunodeficiency, Centromere region instability and Facial anomalies syndrome (ICF) patient cells, abnormally high level of TERRAs have been suggested to give rise to R-loops that cause telomeric dysfunction and genome instability [88].

## Conclusions

Collectively, the work presented here suggest for a model (Figure 4) where HP1γ protects the telomeric chromatin state, through the regulation of telomere and telomere-accessory factors including TRF1 and TERRAs. Loss of HP1γ results in a deprotection of telomeres, uncontrolled expression of TERRAs and accumulation of telomeric R-loops. The lower levels of TRF1, in combination with the elevated R-loops levels cause replication stress which in turn cause DNA damage. The accumulated DNA damage at telomeres can contribute to the onset of senescence, observed in HP1γ^-/-^ cells.

**Figure 4:**
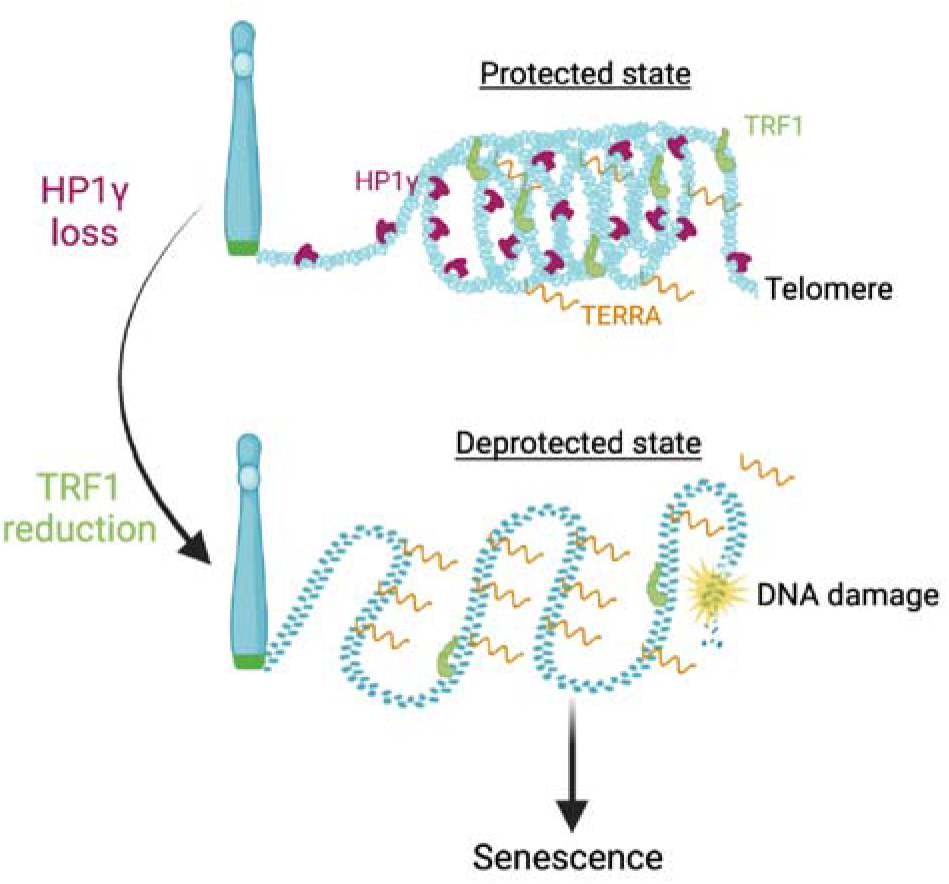
Model of the role of HP1γ on telomere maintenance. Illustration of the protective role of HP1γ on telomere stability. Loss of HP1γ results in a decrease of TRF1 levels. In combination with upregulation of TERRA and formation of telomeric DNA:RNA hybrids, telomeres undergo replication stress resulting in DNA damage and compromise of their integrity. Failure to maintain telomeric chromatin state can lead to earlier onset of senescence.

## CRediT authorship contribution statement

JBV and RF conceptualised the study, designed the experiments and drafted the manuscript. ES performed all the experiments, analysed the data and participated in the manuscript redaction.

## Acknowledgements

JBV is supported by the London Institute of Medical Sciences (LMS), which receives its core funding from UKRI (MRC) and by an ERC Starter Grant (637798; MetDNASecStr) and is thankful to Imperial College London for its support. MS is funded by an MRC PhD fellowship.

## Figure captions

**Figure S1:**
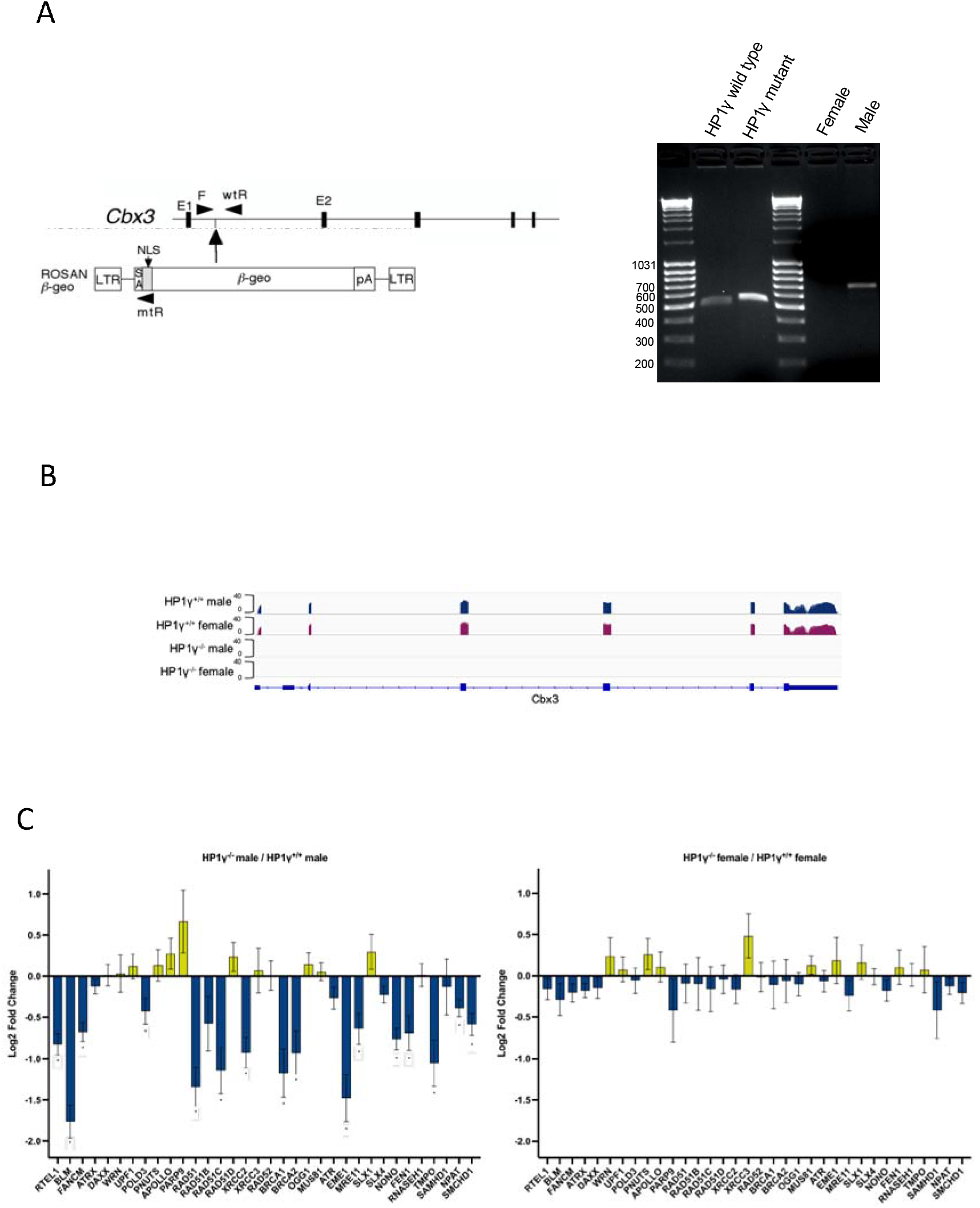
(A) Left: Schematic diagram of the insertion site of the gene trap in the *Cbx3* gene and localisation of primers for genotyping strategy (black arrows), adapted from [34]. Right: Genotyping of E13.5 embryos. PCR for HP1γ genotyping shows a 501bp band for the wild-type allele and a 525bp band for mutant allele, while PCR for Kdm5d genotyping is used for sex determination of embryos. Male animals show a 597bp band while female animals show no bands. (B) Integrative Genome Viewer snapshot of the Cbx3 gene with exons (blue box). The height of each track is proportional to the expression level (log scale) and is the overlay of three biological replicates (analysis from RNA-sequencing available in Preprint BioRxiv: https://doi.org/10.1101/563940) (C) Changes in expression of factors that are necessary for telomere stability upon HP1γ depletion. Bars represent the mean log2 fold changes ± SEM of three biological replicates for each genotype in both sexes. Data was extracted from available Preprint BioRxiv: https://doi.org/10.1101/563940. * padj < 0.05.

**Figure S2:**
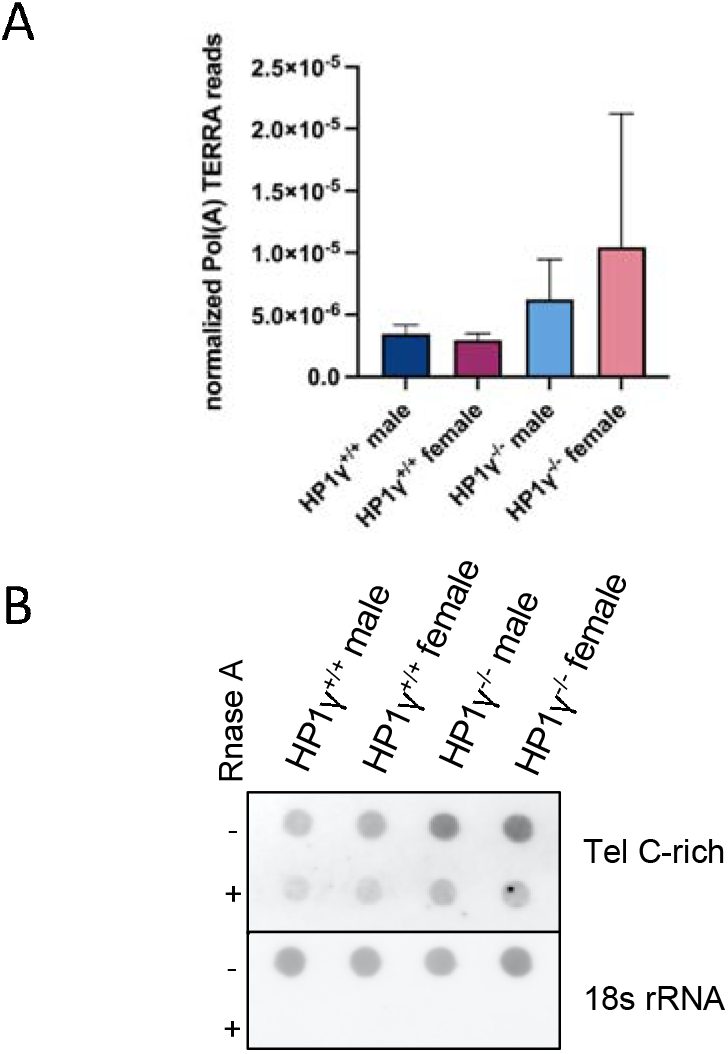
(A) Depletion of HP1γ leads to increased pol(A) transcripts, especially in females. Bars represent the mean of pol(A) TERRA normalized read counts of three biological replicates, from RNA-sequencing data available in Preprint BioRxiv: https://doi.org/10.1101/563940. Error bars indicate SD. No statistical significance as tested with one-way ANOVA and Tukey’s *post-hoc* test. (B) HP1γ loss leads to elevated RNA containing telomeric repeats levels. RNA dot blot analysis of total MEFs RNA (2 μg). The blot was revealed with a DIG-Tel-C-rich probe and an 18 s rRNA probe, which served as a loading control. RnaseA treated samples serve a specificity quality control.

**Figure S3:**
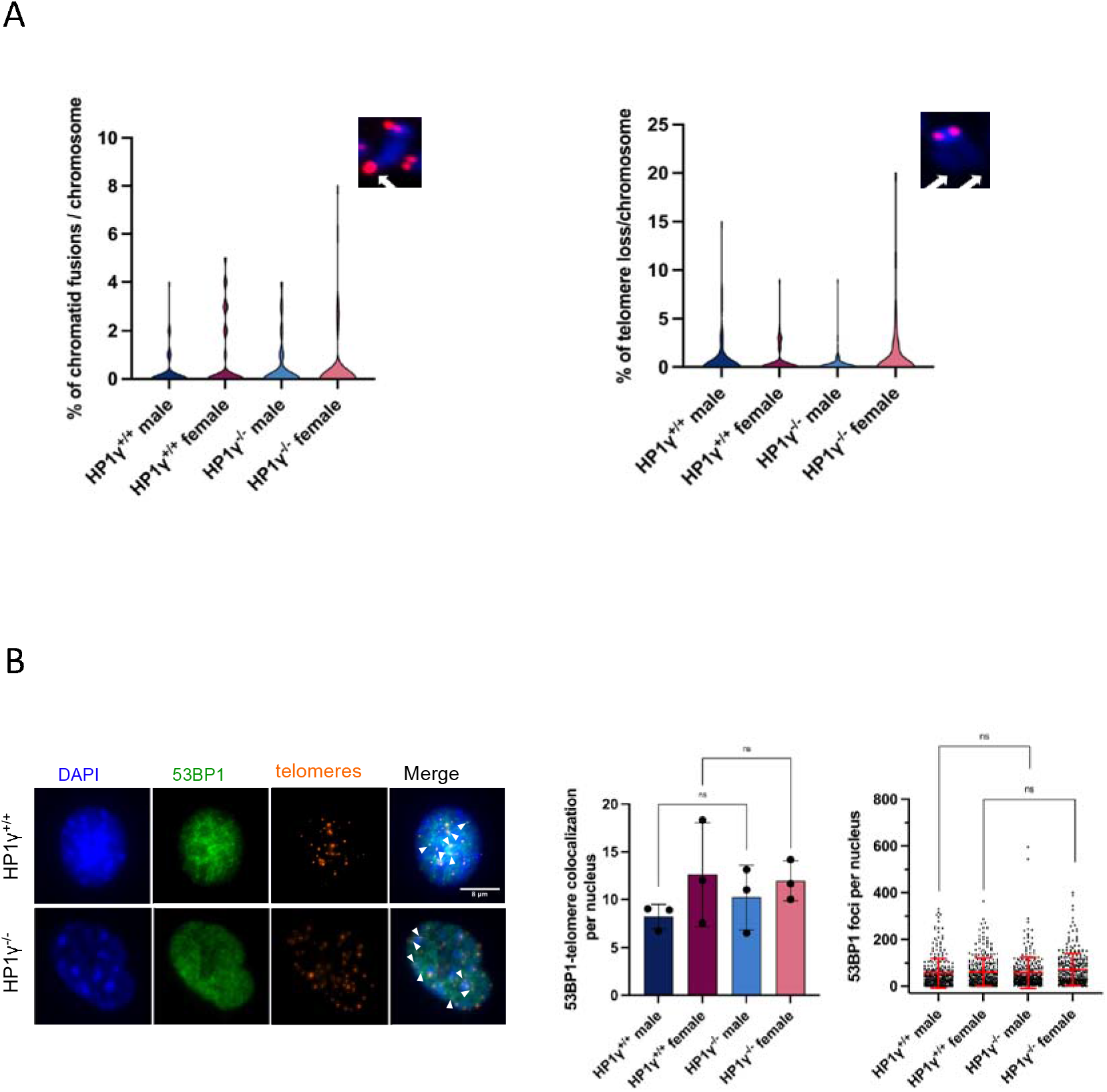
(A) Left: Percentage of chromosome fusions per chromosome. Violin plots represent the combined analysis of 60 metaphases from three independent biological replicates (20 metaphases per biological replicate). Median is coloured in red and quartiles in white. The arrow indicates signal with double the normal intensity (representative sister chromatid fusion site). Right: Percentage of telomere loss per chromosome. Violin plots represent the combined analysis of 60 metaphases from three independent biological replicates (20 metaphases per biological replicate). Median is coloured in red and quartiles in white. The arrow indicates absence of signal (representative telomere loss site). B) Left: Representative image of Immunofluorescence-FISH showing co-localisation of Telomeres (red) with 53BP1 (green) in HP1γ^+/+^ and HP1γ^-/^ MEFs nuclei (DAPI, blue). Middle: Quantification of 53BP1 co-localisation to telomeres per nucleus (n = 300 nuclei). Data are represented as mean ± SD from three independent biological replicates Right: Quantification of 53BP1 foci per nucleus (n = 300 nuclei). Data are represented as mean ± SD from three independent biological replicates

**Table S1.**
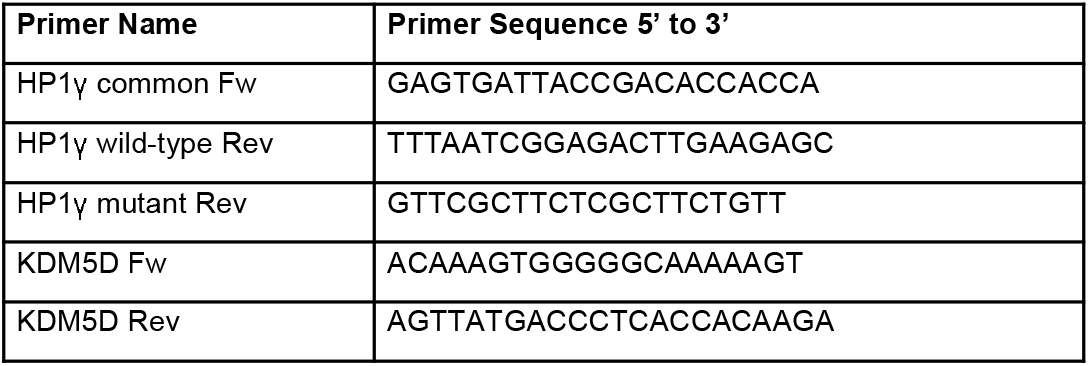
PCR primers used in this study

**Table S2.** HP1γ PCR parameters

| Cycles | Temperature (°C) | Time |
| --- | --- | --- |
| 1 | 96 | 3 min |
| 5 | 96<br>70 (-1 °C per cycle)<br>72 | 15 sec<br>15sec<br>40 sec |
| 30 | 96<br>58<br>72 | 15 sec<br>15 sec<br>45 sec |
| 1 | 72 | 10 min |
| 1 | 12 | hold |

**Table S3.** HP1γ KDM5D parameters

| Cycles | Temperature (°C) | Time |
| --- | --- | --- |
| 1 | 96 | 5 min |
| 35 | 96<br>60<br>72 | 30 sec<br>30 sec<br>15 sec |
| 1 | 72 | 10 min |
| 1 | 12 | hold |

**Table S4.**
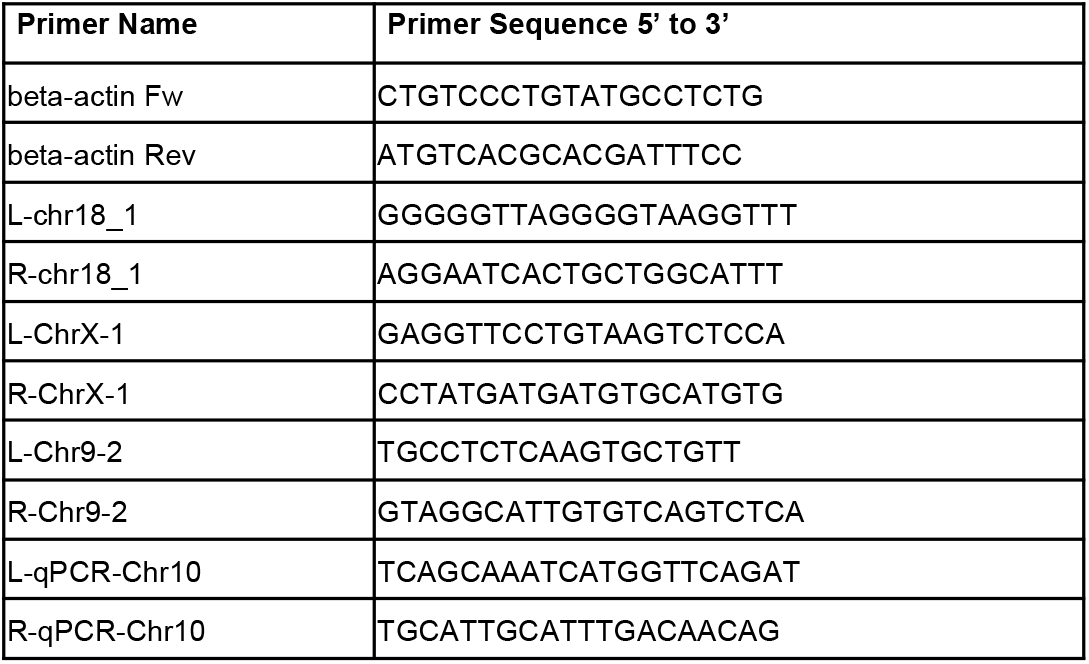
qPCR primers used in this study

